# Molecular and structural basis of a subfamily of PrfH rescuing both the damaged and intact ribosomes stalled in translation

**DOI:** 10.1101/2025.01.09.632186

**Authors:** Yannan Tian, Qingrong Li, Shirin Fatma, Junyi Jiang, Hong Jin, Fuxing Zeng, Raven H. Huang

## Abstract

In bacteria, spontaneous mRNAs degradation and ribotoxin-induced RNA damage are two main biological events that lead to the stall of protein translation. The ubiquitous trans-translation system as well as several alternative rescue factors (Arfs) are responsible for rescuing the stalled ribosomes caused by truncated mRNAs that lack the stop codons. To date, protein release factor homolog (PrfH) is the only factor known to rescue the stalled ribosome damaged by ribotoxins. Here we show that a subfamily of PrfH, exemplified by PrfH from *Capnocytophaga gingivalis* (*Cg*PrfH), rescues both types of stalled ribosomes described above. Our *in vitro* biochemical assays demonstrate that *Cg*PrfH hydrolyzes the peptides attached to P-site tRNAs when in complex with both the damaged and intact ribosomes. Two cryo-EM structures of *Cg*PrfH in complex with the damaged and intact 70S ribosomes revealed that *Cg*PrfH employs two different regions of the protein to recognize two different stalled ribosomes to orient the GGQ motif for peptide hydrolysis. Thus, using a combination of bioinformatic, biochemical, and structural characterization described here, we have uncovered a family of ribosome rescue factors that possesses dual activities to resolve two distinct stalled protein translation in bacteria.

## Introduction

Ribosomal quality control is essential for the health and survival of all organisms. In bacteria, the most frequent impediment of protein translation is the truncated mRNAs that lack stop codons (no-stop mRNAs). Translation of such mRNAs result in the stall of translating ribosomes, and bacteria employ various ribosome rescue factors to resolve the stalling. The ubiquitous trans-translation system is the primary ribosome rescue system, mediated by transfer-messenger RNA (tmRNA) and small protein B (SmpB) (*1–5*). In addition, the alternative rescue factors (Arfs), such as ArfA, ArfB, and ArfT, are also involved in rescuing the non-stop translation in an event that the trans-translation system fails (*6–15*).

In addition to stalled protein translation caused by no-stop mRNAs, damage of essential RNAs for protein translation by ribotoxins also result in ribosome stalling. A complete damage of a particular tRNA by a ribotoxin makes the tRNA unavailable for protein translation, resulting in a stalled ribosome with the empty A-site. In such a case, the only solution appears to be the repair of the damaged tRNA, which will allow the stalled translation to resume.

In addition to many ribotoxins that damage tRNAs, the ribosome is also the target of several ribotoxins. Specifically, rRNA damage at highly conserved sites, such as the decoding center and the sarcin-ricin loop (*16–19*), disable the activity of the ribosome, resulting in ribosome stalling. If the damage site is inaccessible for repair by a RNA repair enzyme, rescuing and dismantling the damaged ribosome is needed, and the damaged ribosomal subunit after separation can then be repaired by an RNA repair enzyme. We have recently demonstrated that two *E. coli* proteins encoded in *rtcB2*-*prfH* operon perform such sequential ribosome rescue and repair events described above (*20*), with *E. coli* PrfH (*Ec*PrfH) being responsible for the rescue. We have carried out extensive biochemical and structural characterization of *Ec*PrfH, providing insight into how *Ec*PrfH specifically recognizes the damaged 70S ribosome for the rescue (*20*).

Upon further bioinformatic analysis of PrfH protein family, we discovered that PrfH can be further classified into two subfamilies based on whether a PrfH possesses an additional C-terminal tail. PrfH of the first subfamily, exemplified by *Ec*PrfH, lack the C-terminal tail. On the other hand, the majority of PrfH from *Bacteroidota* bacteria, exemplified by PrfH from *Capnocytophaga gingivalis* (*Cg*PrfH), possesses a highly conserved C-terminal tail. Here we report our comprehensive biochemical and structural characterization of *Cg*PrfH engaged in rescuing two distinct stalled ribosomes.

## Results

### Bioinformatic analysis of PrfH protein family IPR017509

During our early stage of investigation to elucidate biological functions of PrfH-RtcB pair, frequently encoded in bacterial operons, we employed recombinant proteins from both *Escherichia coli* and *Capnocytophaga gingivalis*. Amino acid sequence alignment of *Ec*PrfH and *Cg*PrfH revealed that, while they are highly homologous in the region encompassing the entire *Ec*PrfH (Extended Data Fig. 1a), *Cg*PrfH possesses an additional C-terminal tail (Fig. 1a). This led us to perform comprehensive bioinformatic analysis of all proteins of IPR017509, which represents PrfH protein family.

**Fig. 1.**
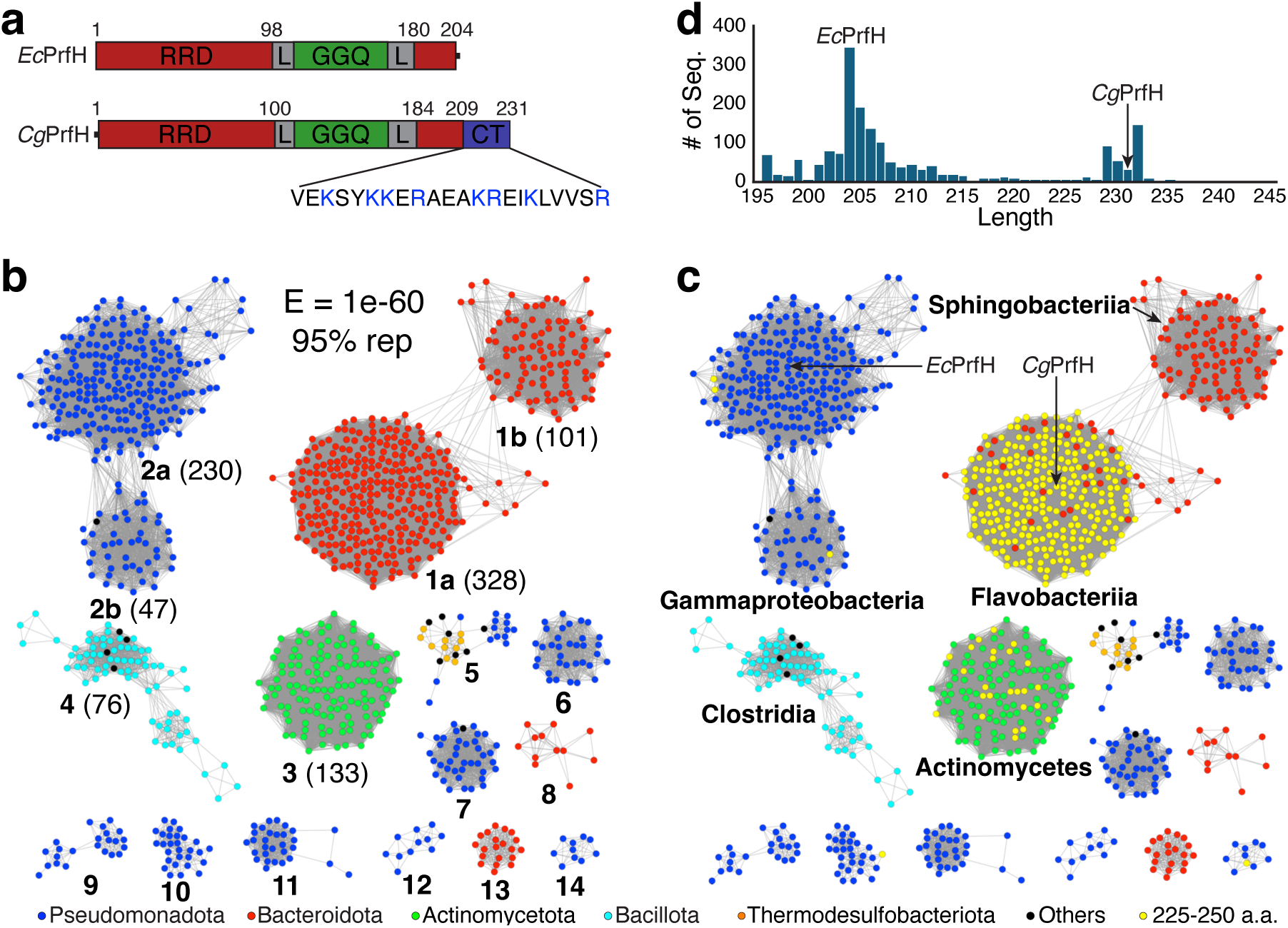
Bioinformatic analysis of PrfH. **a**, Schematic view of the domain structures of *Ec*PrfH and *Cg*PrfH, respectively. As described in our previous study, *Ec*PrfH consists of the ribosome recognition domain (RRD, colored red) and GGQ domain (color green), connected by two linkers (L, colored gray). *Cg*PrfH has the similar domain structures but possesses an additional C-terminal tail (CT, and colored blue), which is rich in basic residues (K and R, colored blue). **b**, Sequence Similarity Network (SSN) of PrfH. Each node (colored circle) represents a collection of PrfH proteins sharing >95% sequence identity (95% rep node). An edge (gray line) connects two nodes if the E-value measuring their sequence similarities is smaller than the cutoff value (1e-60 for this network). The clusters are colored by phyla as indicated at the bottom of the figure. The major clusters are labeled with bold numbers, whose order is based on the total numbers of the nodes within each cluster, which are indicated in parenthesis. **c**, The same SSN as in **b** except the nodes that represent larger PrfH (225-250 a.a.) are highlighted in yellow. Here, the major clusters are labeled by the classes of bacteria. The nodes representing *Ec*PrfH and *Cg*PrfH are marked with arrows. **d**, The histogram of PrfH sizes, which was calculated based on a SSN of 100% rep (representing all PrfH of unique sequences).

We constructed a Sequence Similarity Network (SSN) of PrfH (Fig. 1b). SSN revealed that PrfH is mainly found in five phyla of bacteria (Fig. 1b, colored blue, red, green, cyan, and orange, respectively). Furthermore, our bioinformatic analysis revealed that PrfH is highly conserved. This is supported by a perfect Convergence Ratio score (1.0) of SSN (*21*), very low E value required for reasonable separation of PrfH in SNN (Fig. 1b), and a narrow range of protein sizes (Fig. 1d). This is also supported by comparison of protein sequences between *Ec*PrfH with *Cg*PrfH. Although *E. coli* and *C. gingivalis* belong to different phyla of bacteria (Fig. 1c), PrfH from these two organisms share 37% sequence identities (Extended Data Fig. 1a).

Despite highly conservation of PrfH described above, however, the histogram of PrfH sizes exhibits a bimodal distribution, with *Ec*PrfH and *Cg*PrfH at or near the centers of the two peaks (Fig. 1d). Sequence alignment of *Ec*PrfH and *Cg*PrfH revealed that the size difference of those two PrfH is due to additional C-terminal tail only present in *Cg*PrfH (Extended Data Fig. 1a), suggesting that those two PrfH might have distinct biological functions.

We have previously characterized the biological function of *Ec*PrfH (*20*), which represents the smaller PrfH in the histogram (Fig. 1d). Here we focus our analysis on the larger PrfH represented by *Cg*PrfH (Fig. 1d). To understand the distribution of larger PrfH, we searched for PrfH with sizes in the range of 225-250 a.a. within SSN. The search revealed that the larger PrfH are mainly present in two clusters (Fig. 1c, colored yellow). The overwhelming majority of larger PrfH (261 nodes, 352 organisms) is found in Cluster 1a of SSN, which are bacteria of phylum *Bacteroidota* and class *Flavobacteriia* (Fig. 1c). A small number of larger PrfH (20 nodes, 20 organisms) is also found in Cluster 3, which are bacteria of phylum *Actinomycetota* and class *Actinomycetes*. Further sequential and structural analysis of those larger PrfH in cluster 1a confirmed the presence of additional C-terminal tail like the one observed in *Cg*PrfH (Extended Data Fig. 1b), which is predicted to form a α-helix by AlphaFold2 (*22*). Five additional nodes of larger PrfH that meet the search criteria were also found in other clusters (Fig. 1c). Individual inspection of those five PrfH indicates that they do not possess the additional C-terminal tails found in *Cg*PrfH. Therefore, the larger PrfH possessing a highly conserved C-terminal tail are only present in most *Flavobacteriia* and some *Actinomycetes* bacteria (Fig. 1c). To facilitate discussion, we named PrfH possessing an additional C-terminal tail as PrfH-CT. Our analysis indicates that PrfH-CT constitutes ∼30% of total PrfH. The specific presence of PrfH-CT within *Flavobacteriia* and *Actinomycetes* bacteria suggests that they might have a biological function beyond the one found in smaller PrfH such as *Ec*PrfH. Here we present our comprehensive biochemical and structural characterization of *Cg*PrfH, which represents the clade of bacteria that possess more than 90% of PrfH-CT (Fig. 1c).

### *In vitro* reconstitution of the peptide release activities of *Cg*PrfH

Using similar approaches as in our study of *Ec*PrfH (*20*), we carried out peptide release assays of *Cg*PrfH. Two different ribosome substrates were employed for the assays. One is *E. coli* 70S ribosome specifically damaged in the decoding center. The second is the intact *E. coli* 70S ribosome. Like *Ec*PrfH, *Cg*PrfH can rescue the damaged 70S ribosome (Fig. 2b, green curve). Furthermore, when mRNAs of different lengths were employed, *Cg*PrfH showed approximately similar activities (Fig. 2c), indicating that the activity of *Cg*PrfH is independent of the lengths of mRNAs.

**Fig. 2.**
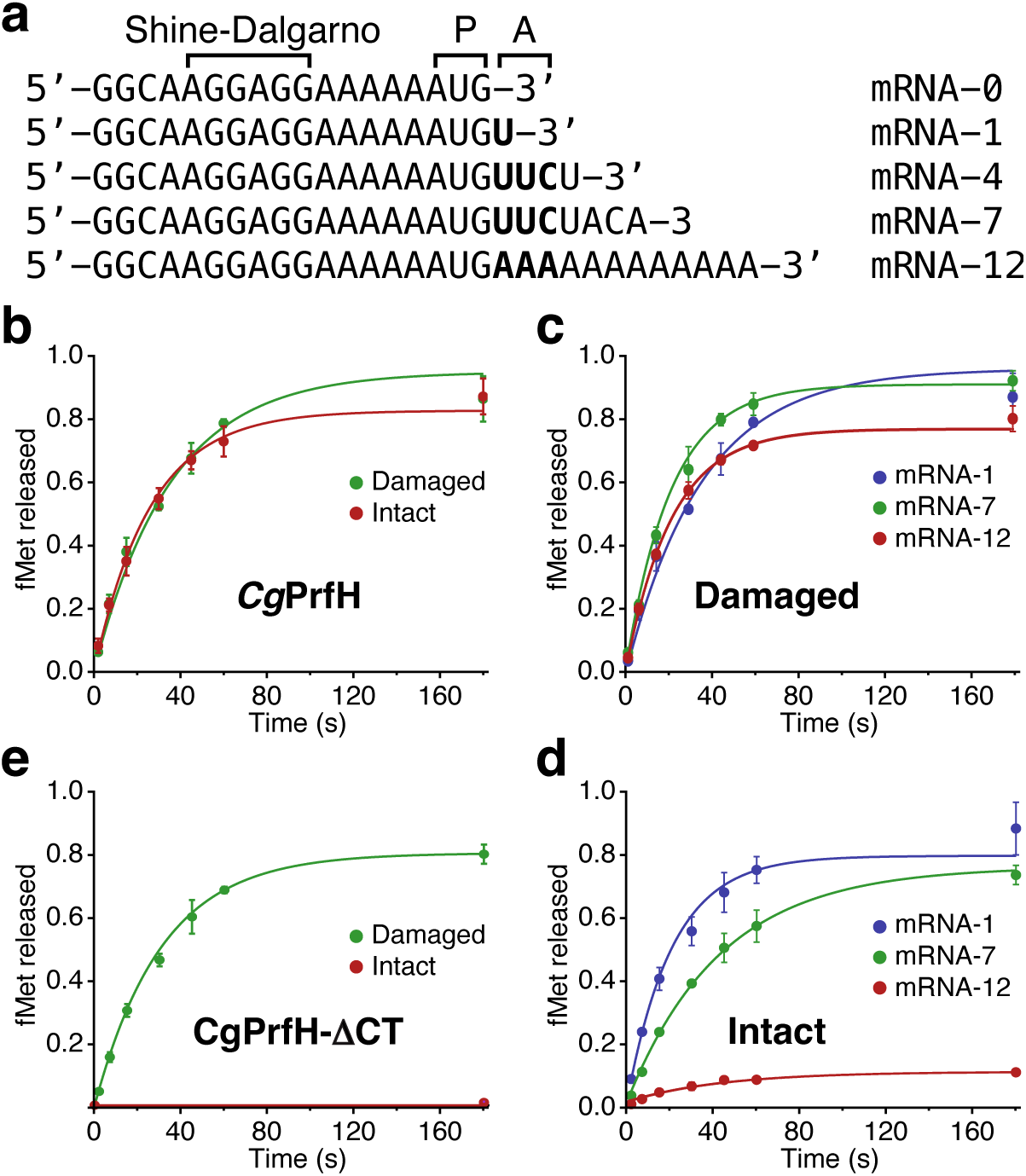
Substrate specificity of peptide release by *Cg*PrfH. **a**, Sequences of five mRNAs employed for this study. P, P-site of ribosome; A, A-site of ribosome. **b**, The time course of the activities of *Cg*PrfH using the damaged 70S ribosome (green) and the intact 70S ribosome (red) as substrates. mRNA-1 was used for the assays. **c**, The time course of the activities of *Cg*PrfH with the damaged ribosome complexed with different lengths of mRNAs. **d**, The time course of the activities of *Cg*PrfH with the intact ribosome complexed with different lengths of mRNAs. **e**, Rescue of damaged and intact ribosomes by *Cg*PrfH-ι1CT, with the C-terminal tail of *Cg*PrfH deleted.

When the intact ribosome in complex with a short mRNA was used as the substrate, *Cg*PrfH is also able to rescue the stalled ribosome with the activity comparable to the one rescuing the damaged ribosome (Fig. 2b, red curve). This contrasts with *Ec*PrfH, as *Ec*PrfH is not able to rescue the stalled intact ribosome (*20*). We also carried out assays with the intact ribosome associated with different lengths of mRNAs, and the rescue activity depends on the length of mRNA (Fig. 2d). Specifically, when a mRNA with 12 nucleotides beyond P-site of ribosome was used, the activity of *Cg*PrfH is greatly diminished (Fig. 2d, red curve). This contrasts with the rescue of the damaged ribosome (compare Fig. 2d to Fig. 2c). Therefore, with the intact ribosome in complex with various lengths of mRNAs, *Cg*PrfH behaves differently from *Ec*PrfH, and acts more like other alternative ribosome rescue factors, ArfB in particular.

To provide further insight into the role of the C-terminal tail in ribosome rescue by *Cg*PrfH, we created a *Cg*PrfH deleting mutant by removing the last 26 residues (named *Cg*PrfH-ΔCT). Peptide release assays revealed that, while *Cg*PrfH-ΔCT maintains its ability of rescuing the damaged ribosome (Fig. 2e, green curve), it is no longer able to rescue the intact ribosome (Fig. 2e, red curve). This result demonstrates that the C-terminal tail of *Cg*PrfH is responsible for the rescue of the intact ribosome.

### Cryo-EM structure of *Cg*PrfH in complex with the damaged 70S *E. coli* ribosome

To provide molecular insight into how *Cg*PrfH rescue the damaged ribosome stalled in translation, we solved the cryo-EM structure of *Cg*PrfH in complex with the damaged *E. coli* 70S ribosome, mRNA-7, and two tRNAs occupying P- and E-sites (Fig. 3 and Extended Data Fig. 2).

**Fig. 3.**
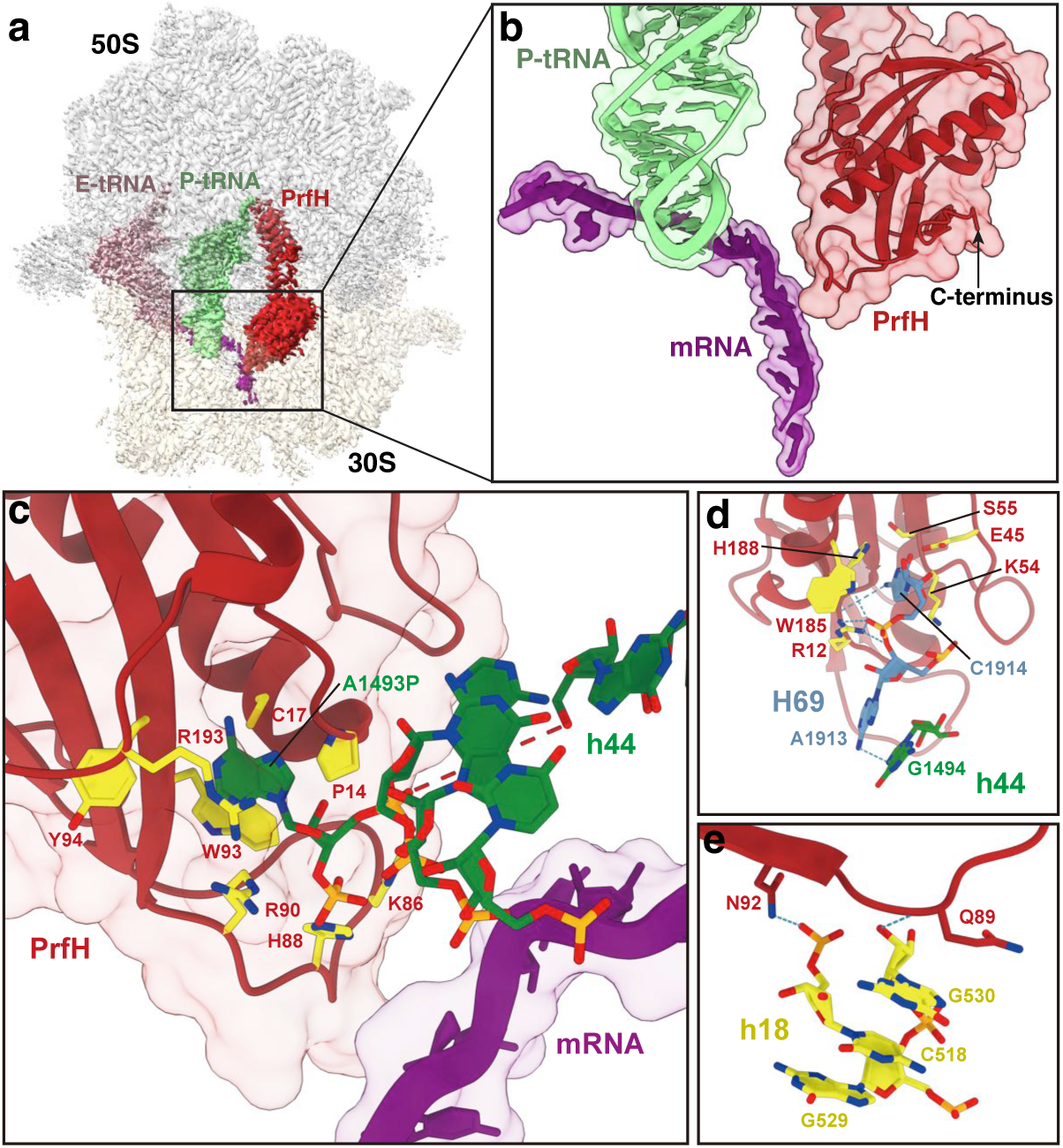
Structure of *Cg*PrfH in complex with the damaged *E. coli* 70S ribosome. **a**, The cryo-EM map of the complex highlighting electron densities corresponding to *Cg*PrfH (red), mRNA (purple), P-site tRNA (light green), and E-site tRNA (light pink). Notice that no electron density corresponding to the C-terminal tail of *Cg*PrfH was observed. **b**, The structure covering RRD of *Cg*PrfH, part of P-tRNA, and the entire mRNA. **c**, Recognition of the 3’-terminal A1493P of the cleaved 16S rRNA by RRD of *Cg*PrfH. **d**, Recognition of C1914 of 23S rRNA and G1494 of 16S rRNA by RRD of *Cg*PrfH. **e**, Specific interactions between RRD of *Cg*PrfH and G530, C518 from h18 of 30S subunit.

Like our previous study of *Ec*PrfH in complex with the damaged 70S ribosome, *Cg*PrfH is found to occupy the empty A-site of the damaged ribosome (Fig. 3a). The cryo-EM structure revealed that, unlike the structural model of *Cg*PrfH via AlphaFold2 (*23*) (Extended Data Fig. 1b), the C-terminal tail of *Cg*PrfH is either unstructured or mobile as the electron density corresponding to the tail was not observed (Fig. 3a,b). Therefore, the structure of *Cg*PrfH in complex with the damaged ribosome is essentially the same as the one of *Ec*PrfH in complex with the damaged 70S ribosome (*20*). Briefly, RRD of *Cg*PrfH specifically recognize the 3’-terminal A1943P of the cleaved 16S RNA

(Fig. 3c). The terminal A1943P is the major determining factor that allows RRD of *Cg*PrfH distinguishing the damaged ribosome from the intact one. In addition, RRD of *Cg*PrfH also makes contacts with nucleotides at the interface of H69 and h44 (Fig. 3d), and it also interacts with G530 and C518 from h18 of 30S subunit (Fig. 3e). Because the structure of *Ec*PrfH in complex with the damaged 70S ribosome has been described extensively (*20*), detailed structural analysis of *Cg*PrfH interacting with the damaged 70S ribosome is omitted here.

### Cryo-EM structure of *Cg*PrfH in complex with the intact 70S *E. coli* ribosome

To provide molecular insight into how *Cg*PrfH rescues the intact ribosome stalled in translation, we solved the cryo-EM structure of *Cg*PrfH in complex with the intact 70S ribosome, mRNA-0 (Fig. 4a), and two tRNAs occupying P- and E-sites (Fig. 4, Extended Data Fig. 3).

**Fig. 4.**
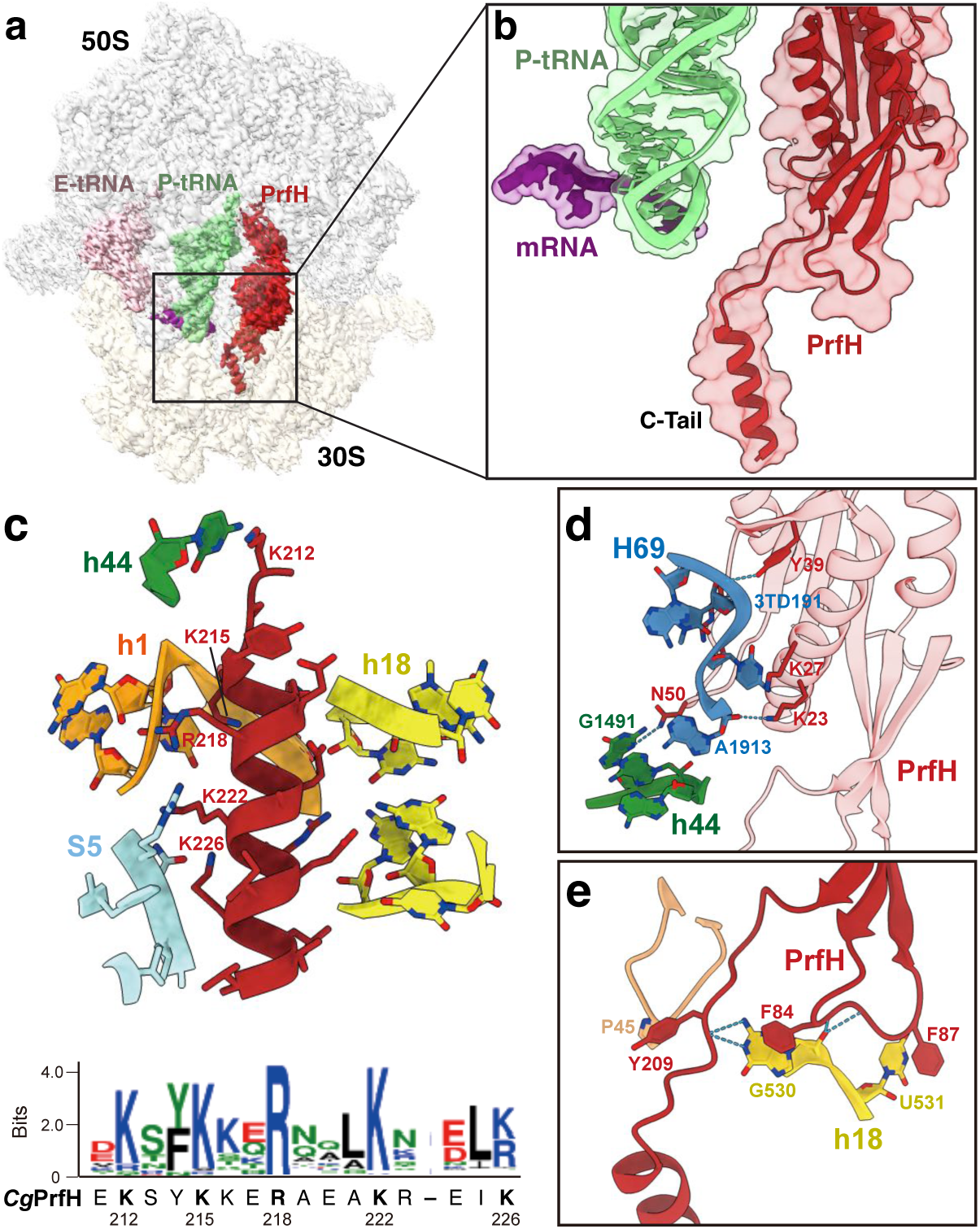
Structure of *Cg*PrfH in complex with the intact *E. coli* 70S ribosome. **a**, The cryo-EM map of the complex highlighting the electron densities corresponding to *Cg*PrfH (red), mRNA (purple), P-site tRNA (light green), and E-site tRNA (light pink). Notice the clear presence of the electron density corresponding to the C-terminal tail of *Cg*PrfH. **b**, The structure covering RRD and the C-terminal tail of *Cg*PrfH, part of P-tRNA, and the entire mRNA. **c**, (Top) Specific interactions between the C-terminal tail of *Cg*PrfH with the wall of the empty mRNA channel. Residues involved in interactions are highlighted with their side chains in stick. Only five highly conserved basic residues are labeled for clarity. (Bottom) HMM of the C-terminal tail of *Cg*PrfH showing high conservation of five basic residues and three hydrophobic residues among larger PrfH. **d**, Specific interactions of RRD of *Cg*PrfH with nucleotides at the interface of H69 of 50S subunit and h44 of 30S subunit. **e**, Additional interactions of RRD of *Cg*PrfH with G530 and U531 from h18 of 30S subunit.

The structure of *Cg*PrfH in complex with the intact ribosome is strikingly different from the one in complex with the damaged ribosome. The most noticeable difference is the C-terminal tail. Unlike the cryo-EM map of the damaged ribosome (Fig. 3a), the electron density corresponding to the C-terminal tail of *Cg*PrfH is clearly present (Fig. 4a), with the helical C-terminal tail occupying the empty mRNA channel (Fig. 4b).

*Cg*PrfH recognizes the intact ribosome through interactions at three locations (Fig. 4c-e). The predominant interactions occur through the C-terminal tail, which makes extensive contacts with the components of the ribosome that form the mRNA channel. The contacts include electrostatic interactions, hydrogen bonding, and hydrophobic interactions. Specifically, seven positively charged residues (K212, K215, K216, R218, K222, R223, and K226) interact with the negatively charged 16S rRNA (Fig. 4c, top). Five of those seven basic residues are highly conserved among PrfH possessing the C-terminal tail (Fig. 4c, bottom). In addition, the side chains of K215, K216, E220, and K222 form hydrogen bonds with three nucleotides of rRNA and one amino acid from ribosomal protein S5. Finally, hydrophobic interactions also occur between several hydrophobic residues in the C-terminal tail of *Cg*PrfH and a hydrophobic patch present by ribosomal protein S5 (Fig. 4c, top).

In addition to the C-terminal tail of *Cg*PrfH, two regions in RRD of *Cg*PrfH that flank the C-terminal tail also make some contacts with the ribosome. One region is residues 23-51 of *Cg*PrfH, making specific interactions with several nucleotides at the interface of H69 and h44 (Fig. 4d). The second region mostly involves a loop consisting of residues 82-89 of *Cg*PrfH, making contacts with G530 and U531 from h18 of 30S subunit (Fig. 4e).

### Understanding the structural and conformational changes in *Cg*PrfH that allow it to rescue two distinct stalled ribosomes

To reveal molecular insight into how a single protein recognizes two different stalled ribosomes for rescue, we performed comprehensive analyses on specific interactions between RRD of *Cg*PrfH and both ribosomes, potential structural changes of individual domains, and structural and conformational changes of the entire *Cg*PrfH protein upon its association with two distinct ribosomes (Fig. 5).

**Fig. 5.**
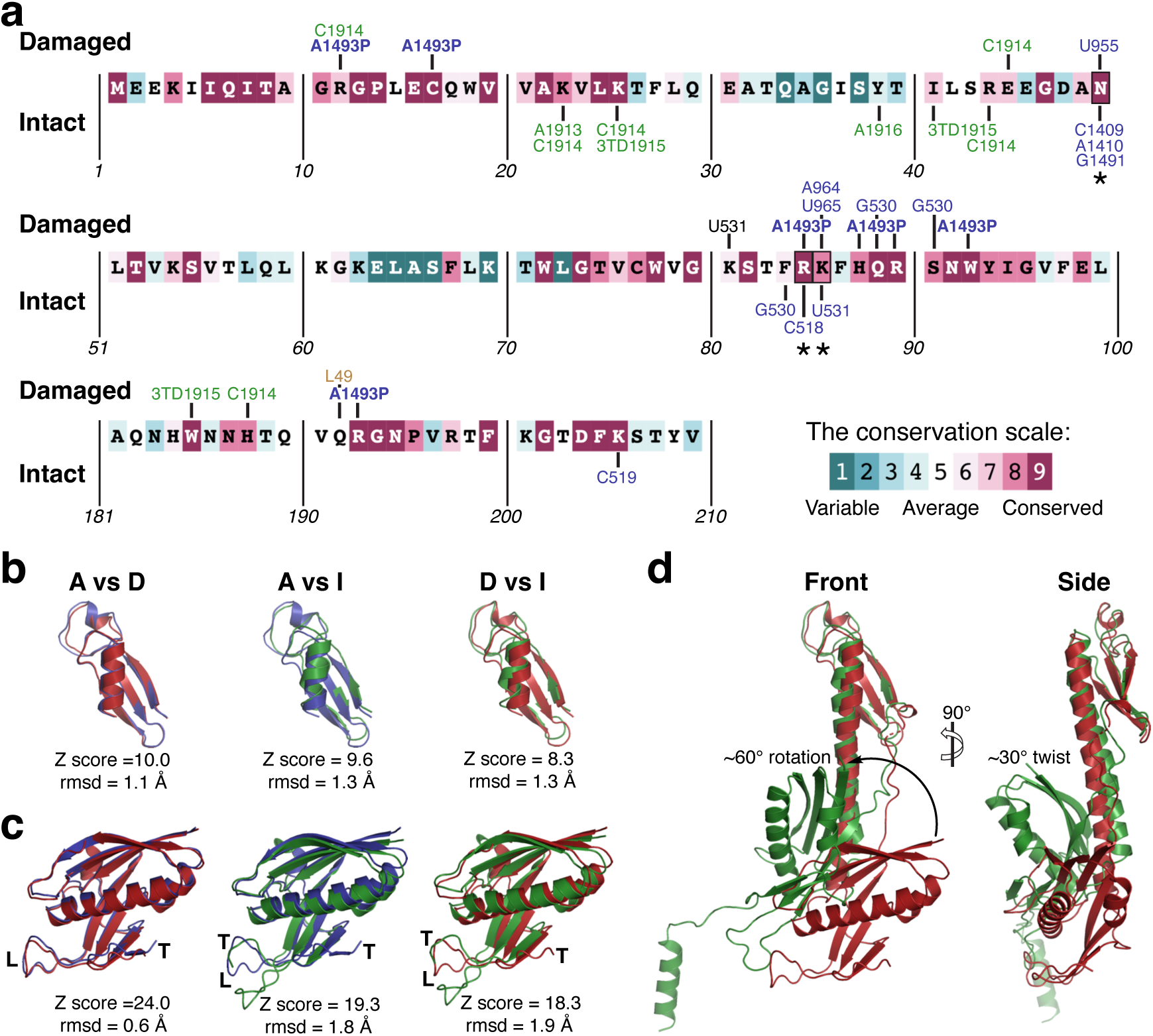
Structural and conformational changes of *Cg*PrfH upon its association with two distinct ribosomes. **a**, Mapping the interactions of RRD of *Cg*PrfH with ribosomes on the amino acid sequence of RRD. The amino acids of RRD of *Cg*PrfH are colored according to their conservations among larger PrfH. The residues from 30S are colored blue, from 50S are colored green, and from ribosomal proteins are colored sand. The 3’-terminal nucleotide A1493P of the cleaved 16S rRNA is highlighted in bold. **b**, Structural alignments of GGQ domain. The structure by Alphafold2 is labeled as A and colored blue, the one bound to the damaged ribosome is labeled as D and colored red, and the one bound to the intact ribosome is labeled as I and colored green. **c**, Structural alignments of RRD. The loops and the peptides leading to the C-terminal tails are labeled as L and T, respectively. **d**, Superposition of the structures of *Cg*PrfH bound to the damaged ribosome to the one bound to the intact ribosome.

We first mapped all contacts between RRD of *Cg*PrfH and two different ribosomes. Thus, the residues within 3.5 Å distance between RRD and the ribosomes were identified and mapped on the sequence of RRD of *Cg*PrfH (Fig. 5a). The amino acids of RRD were colored based on their conservation, which was obtained via ConSurf server (*24*). The analysis identified 16 residues in RRD of *Cg*PrfH that make contacts to the damaged ribosome (Fig. 5a, labels above RRD sequence), and 10 residues that make contacts to the intact ribosome (Fig. 5a, labels below RRD sequence). Only three residues, N50, R85, and K86, make contacts to both ribosomes (Fig. 5a, marked with asterisks). Remarkably, they do not interact with the same nucleotides from two different ribosomes (Fig. 5a). To achieve different interactions described above, RRD of *Cg*PrfH in two different complexes must be structurally and/or conformationally different from one another. In addition, the C-terminal tail of *Cg*PrfH only interacts with the intact ribosome but not the damaged ribosome, which are not present in Fig. 5a.

To provide structural insight into the difference of RRD when interacting with two different ribosomes, we compared the structures of *Cg*PrfH in two complexes together with the one predicted by AlphaFold2 (Fig. 5b-d). We first compared the structures of individual domains. Thus, Dali pairwise structural alignments were carried out for both GGQ domain and RRD. Structural comparisons of GGQ domain revealed that three structures are very similar with each other, with rmsd in the range of 1.1-1.3 Å (Fig. 5b). Structural alignments of RRD also showed that, overall, they are structurally similar (rmsd in the range of 0.6-1.9 Å) (Fig. 5c). However, they have some small but significant differences, which might be functionally important. For example, the structures and conformations of the loop consisting of residue 72-92 are significantly different among the structures compared (Fig. 5c, labeled with L). This is the loop that makes the most contacts with the rRNA-damaged A1493P in the complex with the damaged ribosome (Fig. 5a). Perhaps the most significant difference regarding the recognition of the intact ribosome is the orientation of the segment of peptide immediately preceding the C-terminal tail. The orientation of the peptide in structure I is in opposite direction when compared to ones in the structures A and D (Fig. 5c, labeled as T). This would place the C-terminal tail of *Cg*PrfH at opposite locations when *Cg*PrfH forms complexes with the intact and the damaged ribosome.

Finally, we compared the structures of full-length protein. In this case, we only aligned two *Cg*PrfH structures that are in complex with the ribosomes (Fig. 5d). The alignment revealed that two RRD domains are in completely different positions, and the transformation from one to another requires approximately 60 degrees of rotation and 30 degrees of twist (Fig. 5d). Different positions of RRD in two different complexes explains why different residues in RRD are involved in interaction with ribosomes, and why not a single residue in RRD interacts with the same nucleotide from two different ribosomes (Fig. 5a). Therefore, based on those analyses, we conclude that the different orientations of the C-terminal tail (Fig. 5c), together with the different positions of RRD (Fig. 5d), allows *Cg*PrfH to recognize two different stalled ribosomes for rescue.

### Comparison of *Cg*PrfH to other non-stop ribosome rescue factors

Like *Cg*PrfH in complex with the intact ribosome, many other ribosome rescue factors, such as tmRNA/SmpB, ArfB, ArfA/RF2, and BrfA/RF2, also utilize the empty mRNA channel to anchor GGQ motif for peptide hydrolysis. As shown in Fig. 6, recognition function is achieved by the C-terminal tail of those factors. Most of them adopt a certain degree of α-helical conformation when bound to the ribosome and thereby sense whether the mRNA channel is empty. However, there are some differences in terms of the positions of α-helix as well as the degree of α-helical conformation. Although the α-helix in SmpB is shorter than the one found in *Cg*PrfH, both of them are positioned in approximately the same location in the empty mRNA channel (Compare Fig. 6b to 6c). On the other hand, *Cg*PrfH and ArfB share almost the same length of α-helix, but the positions of α-helices are significantly different (Compare Fig. 6b to 6d). Specifically, the α-helix of *Cg*PrfH is closer to S4 and S5, and distant from h1 and h18, whereas the α-helix of ArfB is less deeply buried in the mRNA channel and closer to the decoding center. Unlike *Cg*PrfH, SmpB, and ArfB, ArfA utilizes unstructured loop instead of α-helix for the recognition of the empty mRNA channel (Fig. 6e). BrfA is more like ArfA except the formation of a very short α-helix at the C-terminal tail (Fig. 6f). Although all those factors adopt different secondary structures to sense the empty mRNA channel, they all employ the positively charged residues for interactions with rRNA surrounding the channel.

**Fig. 6.**
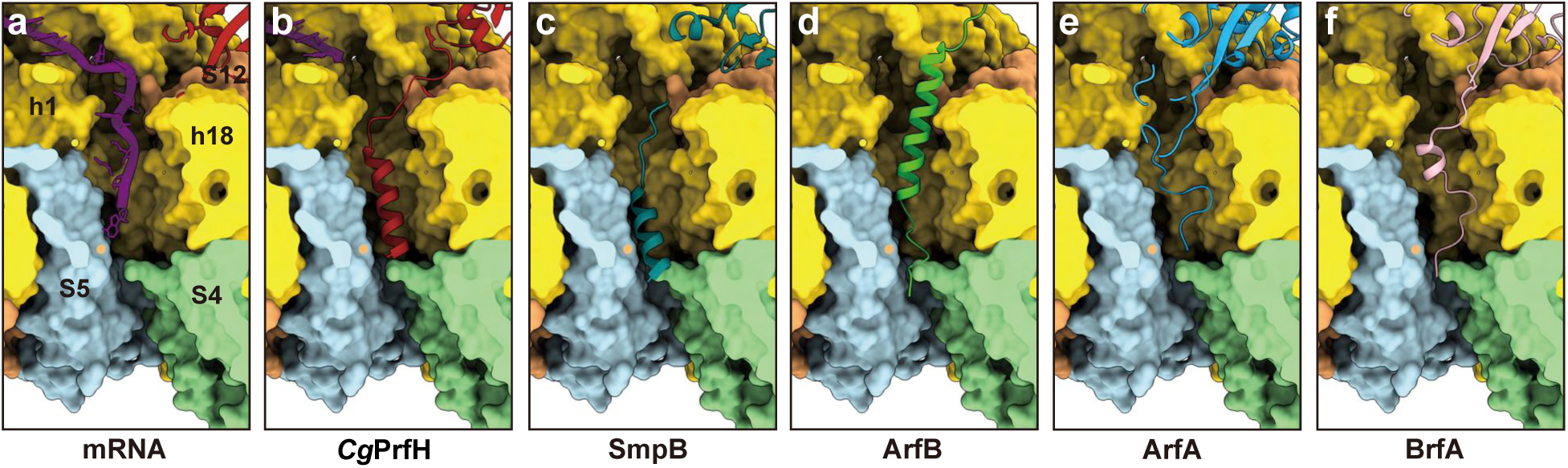
Structural comparison of the C-terminal tails of various non-stop rescue factors inserted into the empty mRNA channel. Cartoon representations of the structures of the C-terminal tails are compared to mRNA as well as to each other. The sources of the aligned structures are: *Cg*PrfH in complex with the damaged ribosome (**a**, this study), *Cg*PrfH in complex with the intact ribosome (**b**, this study), SmpB (**c**, PDB: 4V8Q), ArfB (**d**, PDB: 6YSS) ArfA (**e**, PDB: 5U4I), and BrfA (**f**, PDB, 6SZS).

Based on structural analysis described above, combined with the consideration whether the ribosome rescue factors make additional contacts immediately outside the mRNA channel (Fig. 6, top), one could argue that *Cg*PrfH is mostly resemble to SmpB in its interactions with the surrounding of empty mRNA channel. ArfB is less similar, as it does not make additional contact with the ribosome immediately outside the channel (Fig. 6d). ArfA and BrfA are the most distant from *Cg*PrfH as their C-terminal tails are essentially unstructured.

## Discussion

Based on bioinformatic, biochemical, and structural data presented in this study, we propose a working model of the biological functions of *Cg*PrfH as schematically depicted in Fig. 7.

**Fig. 7.**
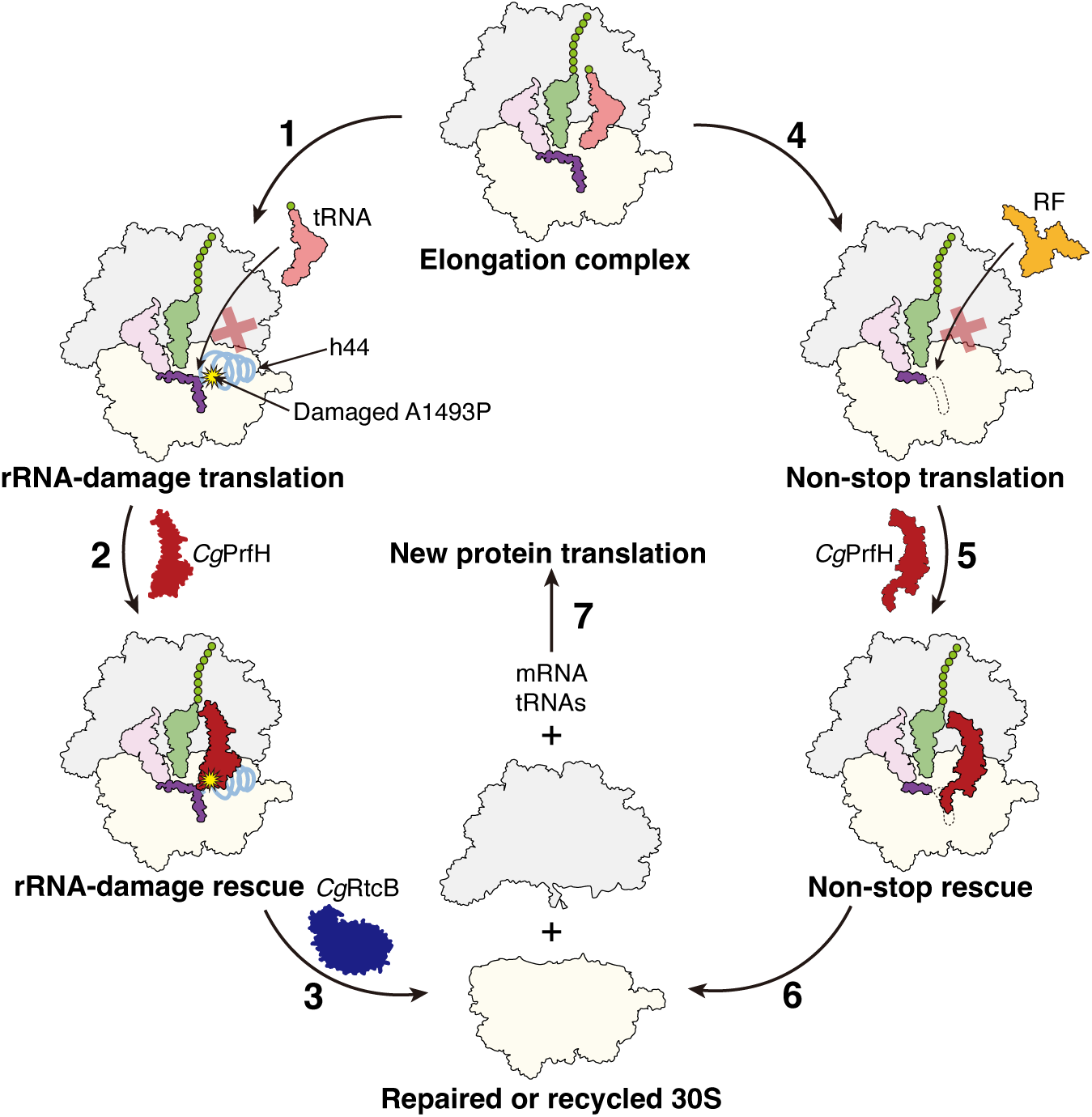
A proposed model of *Cg*PrfH rescuing two different stalled ribosomes. The elongation complex of bacterial protein translation is schematically depicted with the presence of 30S and 50S subunits of the ribosome (colored pale yellow and gray, respectively), three tRNAs occupying A-, P-, and E-sites, and an mRNA occupying the mRNA channel (Fig. 7, top). Additional schematic depictions include h44 of 30S subunit, rRNA-damage site in h44 represented by the 3’-terminal nucleotide A1493P, two *Cg*PrfH with different structures and conformations, and *Cg*RtcB for RNA repair.

An invading ribotoxin such as C-terminal toxin domain of CdiA^ECL^ targets the elongation complex of a translating bacterial 70S ribosome, resulting in specific cleavage of 16S RNA between nucleotides A1493 and G1494 in h44 of 30S subunit. This ribosome damage disables the decoding function of the ribosome, resulting in the failure of delivering a tRNA to A-site and thus ribosome stalling (Fig. 7, step 1). *Cg*PrfH enters the empty A-site of the stalled ribosome and performs ribosome rescue (Fig. 7, step 2). This is mainly made possible by extensive interactions between the 3’-terminal damaged nucleotide A1493P with RRD of *Cg*PrfH. The C-terminal tail of *Cg*PrfH is not involved in recognition of the damaged ribosome. Therefore, the mechanism of rescuing the damaged ribosome by *Cg*PrfH is similar to the one carried out by PrfH that lacks the C-terminal tail such as *Ec*PrfH (*20*). After the hydrolysis of the P-site peptide, the 70S ribosome is dismantled into 30S and 50S subunits, allowing the damaged 30S subunit to be repaired by *Cg*RtcB (Fig. 7, Step 3). The repaired 30S subunit can assemble with a new 50S, a new mRNA, and tRNAs to start a new round of protein translation (Fig. 7, Step 7).

When the elongation complex is translating a truncated mRNA that lacks a stop codon, a RF1 or RF2 cannot enter the A-site near the end of translation due to a lack of a stop codon in mRNA, resulting in a stalled ribosome with an empty mRNA channel (Fig. 7, step 4). *Cg*PrfH is also able to enter the A-site to rescue such a stalled ribosome (Fig. 7, step 5). In this case, the C-terminal tail of *Cg*PrfH anchoring in the empty mRNA channel, not RRD that recognizes the damaged 16S RNA, is mainly responsible for orienting the GGQ motif for the hydrolysis of peptide attached to the P-site tRNA. To achieve this rescue, RRD of *Cg*PrfH adopts a different structure and conformation from the one associated with the damaged ribosome (Fig. 5d). This structural and conformational changes are even apparent with the schematically depiction of *Cg*PrfH shown in Fig. 7. Like the repaired 30S, the rescued 30S from non-stop ribosome can start a new round of protein translation by forming a 70S initiation complex with a new 50S, a new mRNA, and tRNAs (Fig. 7, step 7).

Recently, Feaga and her coworkers performed a comprehensive bioinformatic analysis on non-stop ribosome rescue factors (*25*). Based on 15,259 representative reference genomes, the authors found that approximately 97% bacteria possess tmRNA/SmpB system. Therefore, the trans-translation system is almost ubiquitous in bacteria. On the other hand, ArfA is only found in very limited bacteria, and approximately 58% bacteria possess ArfB. Therefore, not all bacteria have alternative rescue factors based on the study to date. Our search of *C. gingivalis* genome for Arfs revealed that *C. gingivalis* does not have either ArfA or ArfB. The study presented here suggest that, in addition to rescuing stalled ribosome caused by rRNA damage in the decoding center, *Cg*PrfH could play a role of rescuing non-stop ribosome similar to the one carried out by ArfA or ArfB. We acknowledge that we employed a heterogenous system, e.g., ribosome from *E. coli* and PrfH from *C. gingivalis*, for the study described here. Therefore, a more definitive answer regarding the biological functions of *Cg*PrfH other than rescuing the damaged ribosome requires further investigation of *Cg*PrfH in a more homogenous system, such as employing a *Flavobacteriia* bacterium to study *in vivo* function of *Cg*PrfH.

## Methods

### Bioinformatic analysis of IPR017509, the PrfH protein family

Bioinformatic analyses were performed on the database of UniProt 2022_04 and InterPro 91. Calculations were carried out at the EFI website (https://efi.igb.illinois.edu/) (*21*). IPR017509 was submitted for the initial calculation for the Sequence Similarity Network (SSN) of PrfH. After the initial calculation was complete, SSN was finalized with the setting of Alignment Score Threshold = 50 and Minimum Sequence Length = 190. The SSN file with 95% ID, e.g., 95% rep node, was displayed with Cytoscape (*26*) to produce the initial SSN, and yFiles Organic Layout was used for the layout of nodes and edges. Minor adjustments of the positions of a couple of clusters were made to make nodes fit better within the space of the figure, resulting in SSN shown in Fig. 1.

### Cloning, overexpression, and purification of recombinant proteins

The gene encoding *Cg*PrfH (from *Capnocytophaga gingivalis* ATCC 33624) was cloned into pRSF-1 vector, which carries a N-terminal 6xHis tag followed by a SUMO tag. To obtain *Cg*PrfH with the side chain of Glutamine in the GGQ motif methylated, *Cg*PrfH was co-expressed with methyltransferase PrmC in *E. coli* BL21 (DE3) strain at 18 °C overnight induced with 0.5% lactose. Cells were harvested with centrifugation and the pellets were resuspended in lysis buffer (20 mM Tris-HCl, pH 8.0, 500 mM NaCl, 5% glycerol). Cells were lysed using French Press, and cell debris was removed by centrifugation at 20,000 g for 40 min at 4 °C. The supernatant was filtered with a 0.45 μm filter, and the filtered solution was loaded into a HisTrap column (GE Healthcare). The proteins were eluted with the imidazole gradient. The fractions containing the SUMO-tagged PrfH were combined and incubated with Ulp1 protease to cleave the SUMO tag. The untagged *Cg*PrfH was obtained by passing through the second HisTrap column. *Cg*PrfH-ΔCT (residues 1-205) was cloned, overexpressed and purified the same way as described above.

*E. coli* IF1, IF2, and IF3 were overexpressed in *E. coli* BL21 (DE3) cells and purified with Ni-NTA column. N-terminal His-tags in these proteins were removed by HRV-3C protease during purifications. *E. coli* Methionine-tRNA synthetase (MetRS) and methionyl-tRNA formyltransferase (FMT) were overexpressed in *E. coli* BL21 (DE3) cells and purified.

### Preparation of *E. coli* 70S ribosome, charged tRNA and release complex

Both the intact and the damaged 70S ribosomes, and tNRA^fMet^ were purified using the method described previously (*27*). Charging of the tRNA and reconstitution of the release complex were achieved as previously described (*20*). The resulting complexes were resuspended in buffer B (50 mM Tris, pH 7.4, 70 mM NH_4_Cl, 30 mM KCl, 10 mM MgCl_2_, and 5 mM β-mercaptoethanol), aliquoted, and stored at −80°C.

### Peptide release assays

The kinetic experiments on peptide release were carried out as previously described with minor modifications (*28*). First, the intact 70S ribosomal complexes or the damaged 70S ribosomal complexes, containing intact or cleaved *E. coli* 70S ribosome, f-[^35^S]-Met-tRNA^fMet^, and one of the five mRNAs shown in Fig. 2a, was assembled as previously described. The complex (50 nM) then reacted with *Cg*PrfH or *Cg*PrfH-ΔCT of various concentrations at 37 °C.

The reactions were quenched by addition of 5% ice-cold trichloroacetic acid at different time points, and the precipitants were removed with centrifugation at 18,000 g for 10 min at 4 °C to separate f-[^35^S]-Met from f-[^35^S]-Met-tRNA^fMet^. The supernatant was recovered, and the released f-[^35^S]-Met was counted in 2 mL of Bio-Safe II Complete Counting Cocktail (Research Products International). The maximum releasable fMet (fMet_Max_) was determined by incubating the 70S ribosomal complexes (50 nM) with 200 μM puromycin (Sigma-Aldrich) at 37 °C for 30 sec. The fraction of f-[^35^S]-Met was determined by the ratio between the released f-[^35^S]-Met from the reaction and fMet_Max_.

### Cryo-EM grid preparation and data collection

The preparation of complex samples followed the same procedure as the *in vitro* complex assembly described above. For the sample of CgPrfH in complex with the damaged ribosome, damaged 70S ribosomes and mRNA-7 were used in the complex assembly. In contrast, intact *E. coli* 70S ribosomes and mRNA-0 were used to assemble the non-stop complex. The complex sample was diluted to a final concentration of 80 nM with Buffer A. Then, 4 μL of the ribosome sample was added to carbon membrane-covered grids (Quantifoil R1.2/1.3, 300 mesh, copper). The grids were quickly blotted and frozen in liquid ethane using a Vitrobot (Thermo Fisher Scientific, USA). Cryo-EM data collection was performed at the Cryo-EM Centre at the Southern University of Science and Technology. A 300 keV Titan Krios electron microscope (Thermo Fisher Scientific, USA) equipped with a Post-GIF Gatan K3 Summit electron detector (Gatan, USA) was used for data collection. All movies were automatically collected using the EPU software (Thermo Fisher Scientific, USA). The data collection parameters included a pixel size of 0.84 Å/pixel, a total electron dose of 30 electrons/Å2, an exposure time of 5 seconds, and a total of 30 frames. Two data sets, one for *Cg*PrfH in complex with the damaged 70S ribosome, and the other for *Cg*PrfH in complex with the intact ribosome, were collected and followed the same data processing pipeline.

### Cryo-EM data processing

Data process was performed using CryoSPARC (*29*). In brief, the micrographs were generated from the raw movies by applying motion correction with MotionCorr2 software (*30*) and CTF estimation with CTFFIND4 (*31*). Particle picking was performed automatically using bod picker and particles were subsequently extracted from micrographs with a box size of 384 pixels. Poorly aligned and contaminant particles were filtered out through 2D classification. Selected particles from the resulting 2D classes were further used for initially modeling. Heterogeneous refinement was performed using the four classes, then remove the trunk particles and poor aligned classes. Density maps containing *Cg*PrfH at the A-site were chosen for homogeneous refinement. CTF refinement was applied to enhance the resolution of the cryo-EM maps.

### Model building and refinement

For model building and refinement, the initial models used were the cryo-EM structure of *Ec*PrfH in complex with the damaged ribosome (PDB: 4SA4) and the Alphafold2-predicted model of *Cg*PrfH. The initial models were fitted into the density map using UCSF Chimera (*32*), and the models of *Cg*PrfH were adjusted based on A-site densities using Coot (*33*). Further refinement was performed in Phenix (*34*). The final models were validated by the MolProbity (*35*), and the structures and maps were visualized using UCSF ChimeraX (*36*) and PyMOL.

### Data availability

Electron microscopy maps have been deposited in the Electron Microscopy Data Bank under accession codes EMD-48479 for the damaged 70S•CgPrfH complex and EMD-48513 for the intact 70S•CgPrfH complex. Coordinates have been deposited in the Protein Data Bank under accession codes 9MOR for the damaged 70S•CgPrfH complex, and 9MQ4 for the intact 70S•CgPrfH complex.

## Acknowledgements

F.Z. is an investigator of SUSTech Institute for Biological Electron Microscopy. We thank the cryo-EM center of Southern University of Science and Technology for cryo-EM data collection, and C. S. Hayes (University of California Santa Barbara) for providing us the plasmids that encodes CdiA-CT^ECL^ and its immunity protein. This work was supported by the National Institute of Health (Grants GM145060 and GM120764 to R.H.H., and Grant GM120552 to H.J.), and by the National Natural Science Foundation of China (Grant 32171200 to F.Z.).

## Author contributions

R.H.H. conceived the study; Y.T., Q.L., F.Z., and R.H.H. designed the experiments; Y.T., Q.L., S.F., and J.J. performed the research; H.J. supervised the early stage of structural study of the intact complex; Y.T., Q.L., F.Z., and R.H.H. analyzed the data; and Y.T., Q.L., F.Z., and R.H.H. wrote the paper with input from other authors.

