## Extended Data Figures and Table for "Molecular and structural basis of a subfamily of PrfH rescuing both the damaged and intact ribosomes stalled in translation"

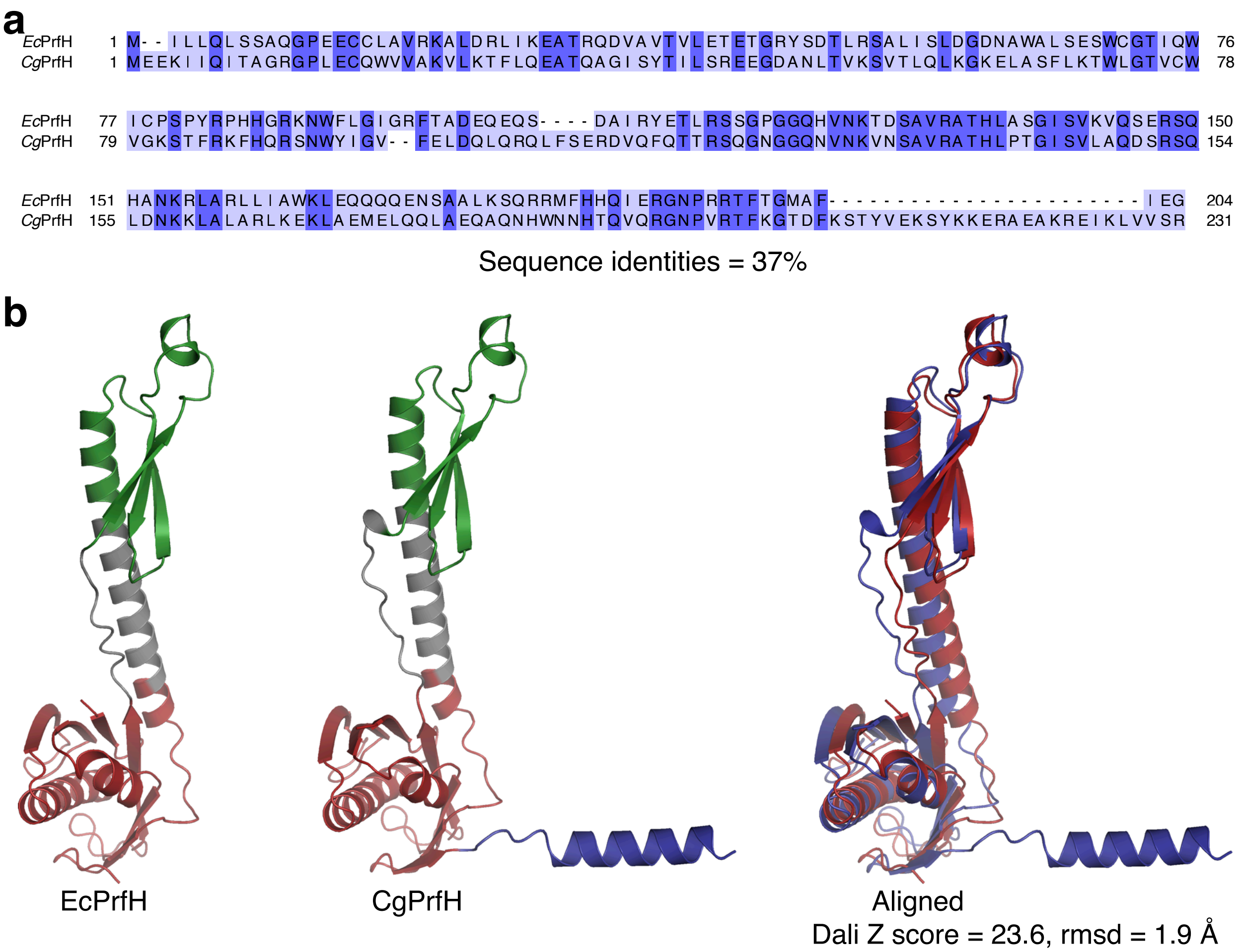


**Extended Data Fig. 1 | Comparing *Cg*PrfH to *Ec*PrfH. a**, Sequence alignment of *Ec*PrfH and *Cg*PrfH. The sequences are colored based on the degree of conservation. **b**, Structural comparison of two PrfH. The structures were obtained via AlphaFold2, and they are colored according to the color scheme in Fig. 1a.

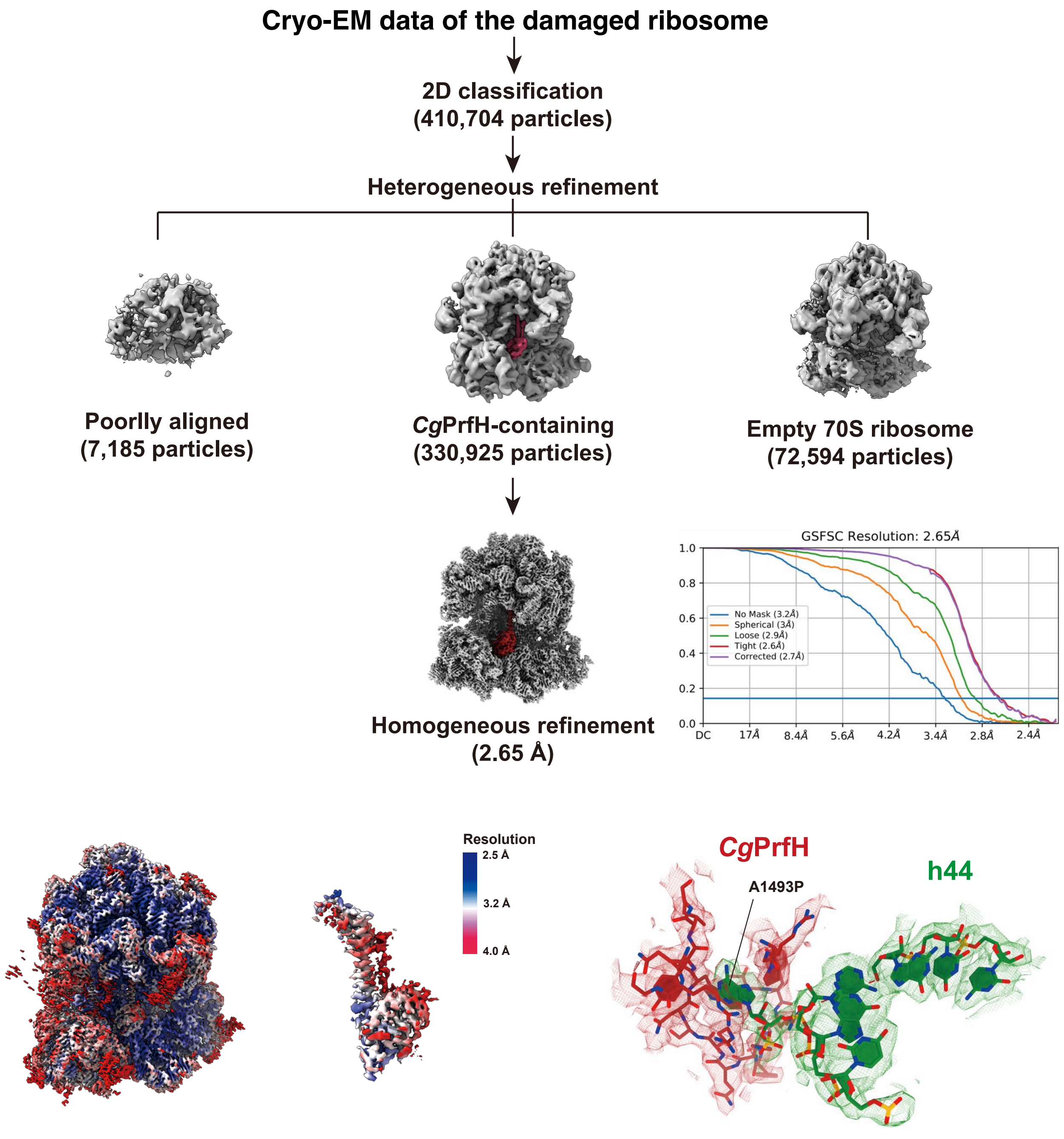


**Extended Data Fig. 2 | Cryo-EM classification of *Cg*PrfH in complex with the damaged *E. coli* 70S ribosome.**

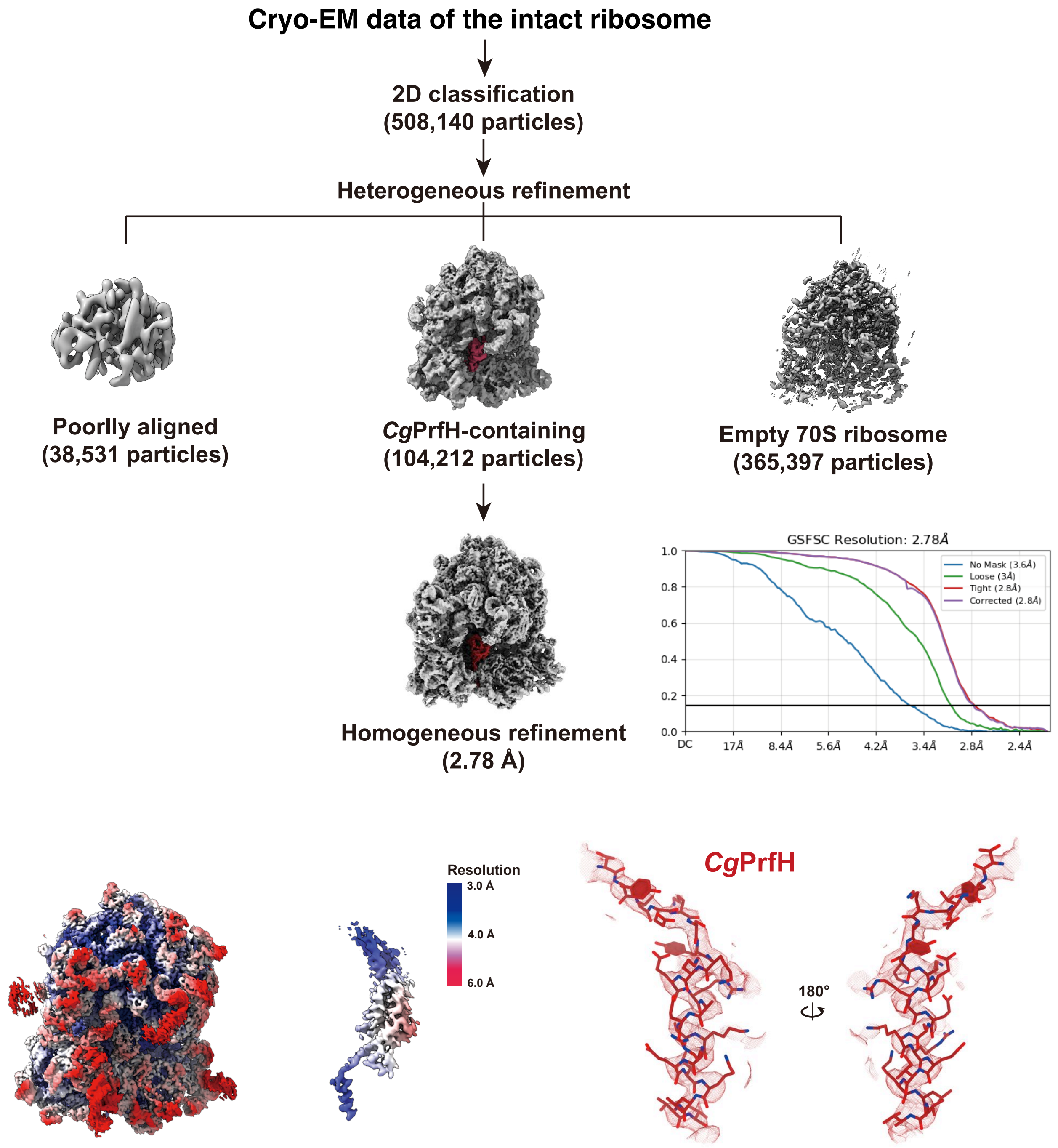


**Extended Data Fig. 3 | Cryo-EM classification of *Cg*PrfH in complex with the intact *E. coli* 70S ribosome.**

**Extended Data Table 1 | The statistics of data collection, refinement, and model**

| | Damaged ribosome complex | Intact ribosome complex |
|---|---|---|
| **PDB entry code** | **9MOR** | **9MQ4** |
| **Data collection and process** | | |
| Camera | Gatan K3 | Gatan K3 |
| Voltage (kV) | 300 | 300 |
| Electron exposure ($e^{-}Å^{-2}$) | 30 | 30 |
| Defocus range (μm) | 1.0~2.5 | 1.0~2.5 |
| Pixel size (Å) | 1.095 | 1.050 |
| Symmetry imposed | C1 | C1 |
| Micrographs collected (no.) | 3,179 | 3,956 |
| Final particle images (no.) | 330,925 | 104,212 |
| Map resolution (Å) | 2.65 | 2.78 |
| **Refinement** | | |
| Model resolution (Å) | 2.7 | 2.8 |
| **Model composition** | | |
| Non-hydrogen atoms | 150,119 | 150,154 |
| Protein residues | 6,180 | 6,201 |
| Nucleotide residues | 4,724 | 4,717 |
| Ligands | 441 | 441 |
| **R.m.s deviations** | | |
| Bond lengths (Å) | 0.33 | 0.85 |
| Bond angles (°) | 0.85 | 0.82 |
| **Validation** | | |
| Molprobity score | 2.11 | 2.11 |
| Clash score | 17.09 | 16.23 |
| Poor rotamers (%) | 0.06 | 0.02 |
| **Ramachandran plot** | | |
| Favored (%) | 94.39% | 94.12% |
| Allowed (%) | 5.62% | 5.81% |
| Disallowed (%) | 0.02% | 0.07% |
